# Reversible translocation of acyl-CoA:cholesterol acyltransferase (ACAT) between the endoplasmic reticulum and vesicular structures

**DOI:** 10.1101/2023.06.06.543826

**Authors:** Andrea Schiffmann, Lena Ahlswede, Gerald Gimpl

## Abstract

The enzyme acyl-CoA:cholesterol acyltransferase (ACAT) is normally localized in the endoplasmic reticulum (ER) where it can esterify cholesterol for storage in lipid droplets and/or the formation of lipoproteins. Here, we report that ACAT can translocate from the ER into vesicular structures in response to different ACAT inhibitors. The translocation was fast (within minutes), reversible and occurred in different cell types. Interestingly, oleic acid was able to fasten the re-translocation from vesicles back into the reticular ER network. The process of ACAT translocation could also be induced by cyclodextrins, cholesterol, lanosterol (but not 4-cholestene-3 one), 25-hydroxycholesterol, and by certain stress stimuli such as hyperosmolarity (sucrose treatment), temperature change, or high-density cultivation. *In vitro* esterification showed that ACAT remains fully active after it has been translocated to vesicles in response to hyperosmotic sucrose treatment of the cells. The translocation process was not accompanied by changes in the electrophoretic mobility of ACAT, even after chemical crosslinking. Interestingly, the protein synthesis inhibitor cycloheximide showed a stimulating effect on ACAT activity and prevented the translocation of ACAT from the ER into vesicles. The translocation process of ACAT may provide a new way to regulate cholesterol esterification in cells, by altering the accessibility of the enzyme to its substrate.

## INTRODUCTION

In cells, the majority (60% - 90%) of total cholesterol is localized in the plasma membrane [28,32,62] where it is organized in different pools [17,33,34,44,47]. Only a small fraction of ‘active’ cholesterol that is not complexed with other lipids (in particular sphingomyelin) can move to the endoplasmic reticulum (ER) by a non-vesicular pathway using transfer proteins of the Aster (= GramD) family [43,53]. As a co-ligand of the Aster proteins, phosphatidylserine also plays a central role in this protein-mediated transfer of cholesterol from the plasma membrane to the ER [20,59,60]. The ER contains only small amounts of total cholesterol and functions as the cholesterol sensing compartment where key components of the cholesterol homeostatic machinery (SREBP-SCAP-INSIG) are localized [6]. Any influx of cholesterol into the ER exceeding a threshold value of ∼5 mol% activates this feedback system. To prevent the accumulation of free cholesterol, which is toxic to cells, cholesterol is esterified by acyl-CoA:cholesterol acyltransferase (ACAT).

ACAT enzymes are also known as sterol O-acyltransferases (SOATs) and exist in two isoforms, ACAT1 and ACAT2. ACAT1/SOAT1 is ubiquitously expressed in nearly all tissues to maintain cholesterol homeostasis, whereas ACAT2 is mainly found in the liver and intestine [51]. They both catalyse the formation of cholesterol esters (CE) from cholesterol and long-chain or medium-chain fatty acids. These esters are either stored in lipid droplets or are secreted as lipoproteins [9,10,36]. ACAT enzymes are membrane proteins residing in the endoplasmic reticulum (ER). However, it is important to mention that a localization of ACAT1 distinct from the ER has also been demonstrated under some conditions and/or in some cell types [29,31,52]. Despite its key role in the cellular sterol homeostasis, the sterol regulation of ACAT1 does not occur at the transcriptional level, as the gene promoter lacks sterol response elements [8,9]. ACATs are allosteric enzymes that form tetramers and can be activated by a variety of sterols. According to one model, ACAT1 contains two different binding sites for steroidal molecules, a substrate and an activator site [50,51]. Recent structural data confirmed the dimerization/tetramerization of ACAT1 [24,46]. Interestingly, ER-resident ACAT1 was shown to esterify also cholesterol metabolites which are produced in the mitochondria such as pregnenolone and some oxysterols [51]. This raises the question of where and how the enzyme interacts with these substrates. Notably, ACAT1 was reported to be enriched in a sub-compartment of the endoplasmic reticulum (ER), the so-called mitochondria-associated membranes (MAMs) [1]. These structures are physically and biochemically connected to mitochondria and could possibly enable direct molecular contacts of enzyme and substrate. But this hypothesis remains to be tested.

In this study, we explored the intracellular distribution profile of ACAT1/SOAT1 under various conditions. Thereby, we observed a significant translocation of the enzyme out of the extensive ER network into vesicular structures in response to certain stimuli. This translocation process was rapid and reversible and could be a way to regulate cholesterol esterification in cells, by altering the accessibility of the enzyme to its substrate. Finally, the results of this study also shed new light on the mechanisms of ACAT inhibitors and may help to explain previous observations regarding the localization profile and the activity of ACAT1/SOAT1.

## RESULTS

### The fluorescent cholesterol analogue DChol and the fatty acid probe Bodipy-C12-568 are reporters of ACAT activity

We have introduced DChol (Fig. 1A) as a valuable reporter of cholesterol trafficking in cells [66]. DChol and cholesterol can easily be incorporated into methyl-β-cyclodextrin (MβCD) cages to form efficient sterol-MβCD donor complexes. This allows rapid sterol delivery to the cells. When CHO cells pulse-labeled with DChol-MβCD were incubated for 24 h in culture medium and then processed for lipid extraction, substantial amounts of DChol were found to be esterified as shown by thin layer chromatography and subsequent fluorescence detection (Fig. 1B). The DChol-ester spot was absent when the cells were treated in the presence of SAN 58-035, an ACAT inhibitor (iACAT) (Fig. 1B). This proves the ACAT specificity of the DChol-ester formation.

**Fig. 1.**
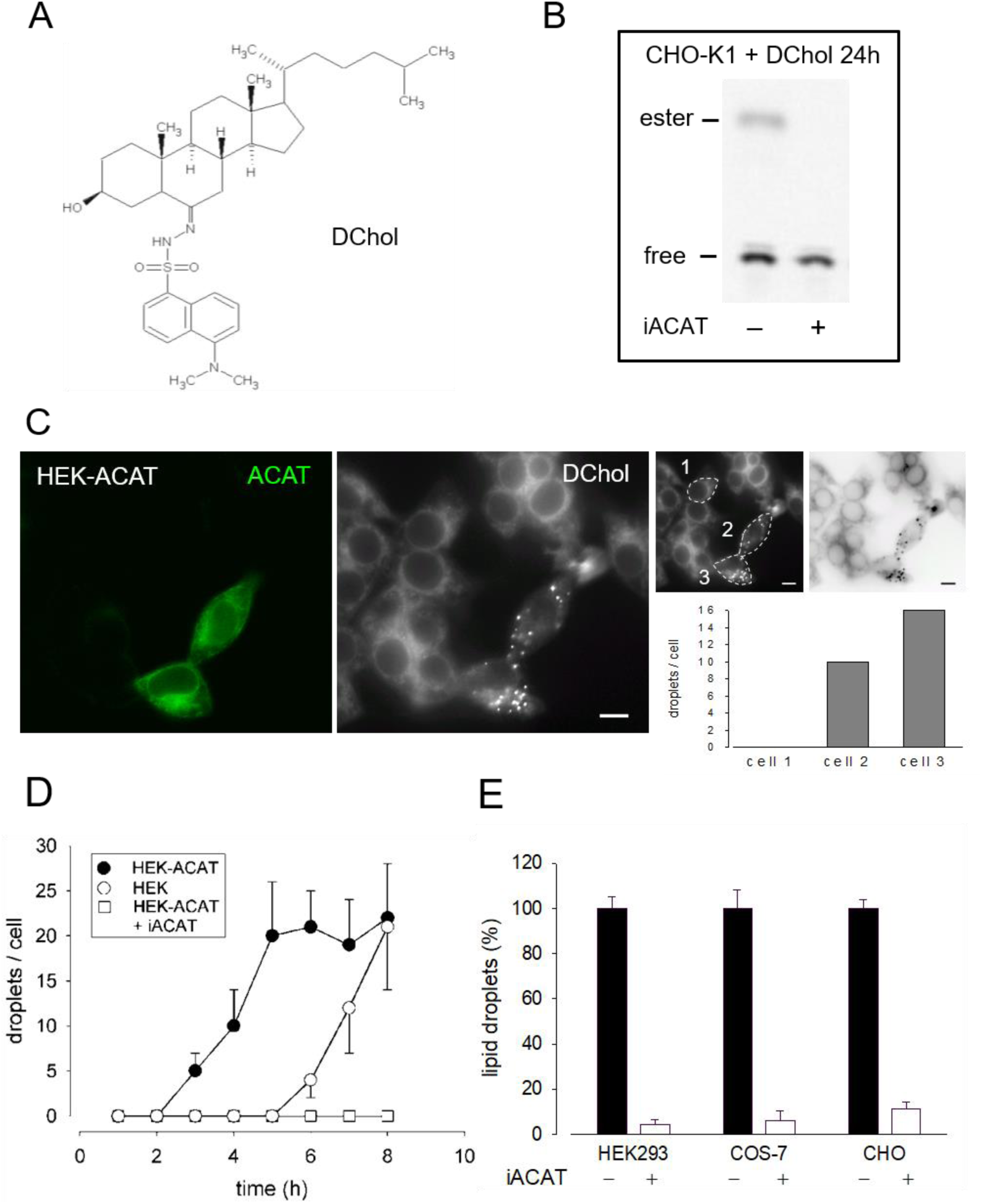
DChol is esterified by ACAT. (A) Structure of DChol. (B) CHO-K1 cells were pulse-labeled for 1 h with DChol-MβCD (7 µM) or Chol-MβCD (7 µM) (’DChol-’) in culture medium in the absence of presence of the ACAT inhibitor SAN 58-035 (iACAT) (0.4 µg/ml). The medium was washed off and chasing was performed for the indicated time. The cells were harvested and their lipids were extracted and separated by silica gel TLC using heptane/diethyl ether/methanol/acetic acid (80:30:3:1.5, by vol) as mobile phase. The fluorescence of free or esterified DChol was detected under UV light. (C) Increase of ACAT activity accelerates the formation of DChol/ester containing lipid droplets. HEK293 cells stably expressing GFP-tagged ACAT1 (HEK-ACAT) were mixed with untransfected HEK293 cells, were co-cultivated for 24 h, and were incubated for 10 min with culture medium containing DChol-MβCD (10 µM). After removal of the medium, the cells were cultivated for 5 h and then observed by fluorescence microscopy. Left image, ACAT; middle image, DChol; right images, quantification of the number of lipid droplets in cell #1 (control), and cells #2 and #3 that overexpress ACAT (bar, 10 µm). (D) HEK cells (ο), HEK-ACAT cells in the absence (□) or presence (•) of iACAT were labeled for 10 min with DChol-MβCD (10 µM). Thereafter, the cells were incubated for the indicated times and the number of lipid droplets were counted (n>50 cells; means ± SD). (E) HEK, COS-7, and CHO cells were labeled for 10 min with DChol-MβCD (10 µM) in the presence or absence of iACAT. After a further incubation of 8 h in culture medium, the DChol containing lipid droplets were counted (n>50 cells; means ± SD).

DChol moves from the plasma membrane to the endoplasmic reticulum (ER) and is then finally transported to lipid droplets where it is stored in form of DChol-ester. This process requires ACAT activity [66]. We therefore expected that overexpression of ACAT should accelerate this storage process. To verify this claim, we conducted an experiment in which HEK293 cells stably expressing EGFP-tagged ACAT1 (termed HEK-ACAT) were mixed with non-transfected HEK293 cells. The cells were pulse-labeled with DChol-MβCD and co-cultivated in medium. After 5 hours of incubation, DChol was mainly found in ER structures in non-transfected cells whereas in cells overexpressing ACAT, lipid droplets containing DChol were clearly present (Fig. 1 C, compare left and middle image). To quantitate this process, DChol-labeled lipid droplets in transfected *versus* non-transfected cells were counted. In the right panel of Fig. 1C, such a quantification is shown for three single cells (compare non-transfected cell #1 with ACAT overexpressing cells #2 and #3). Thus, the development of DChol containing lipid droplets was significantly accelerated in cells that overexpress ACAT. Under the same experimental conditions, the formation of DChol-labled lipid droplets occurred ∼4 h earlier in HEK cells overexpressing ACAT compared with non-transfected HEK cells (Fig. 1D). After an incubation time of 8 h, roughly equal amounts of lipid droplets were visible in non-transfected *versus* ACAT overexpressing HEK cells. In the presence of an ACAT inhibitor, the appearance of lipid droplets was completely suppressed during the time course of the experiment (Fig. 1D). The appearance of ACAT specific DChol-labeled lipid droplets also occurred in other cells. After 8 hours, the formation of DChol containing lipid droplets in dependence on the activity of ACAT was also observed in CHO and in COS-7 cells (Fig. 1E).

The esterification of cholesterol also requires a continuous supply of fatty acids. The incorporation of a fluorescent fatty acid could therefore also report on the formation of DChol esters. For this purpose, we performed labeling experiments with the fatty acid probe Bodipy-C12-568 in AC29 cells which are mutant CHO cells deficient in endogenous ACAT activity. AC29 cells stably transfected with AcGFP-ACAT1 (termed AC29-ACAT cells) were pulse-labeled for 10 min with Bodipy-C12-568. After washing and chasing for 3 h with culture medium, the cells were analyzed by fluorescence microscopy. As shown in Fig. 2, only the three ACAT expressing cells (left upper image) have incorporated substantial amounts of Bodipy-C12-568 (middle upper image and right overlay image). The quantification of these signals in 3 regions of interest (roi) (Fig. 2, lower images: #1, non-transfected cell; #2 and #3, transfected cells) revealed large differences (∼150 fold) in the fluorescence of the fatty acid probe between ACAT expressing and non-transfected cells. Thus, the incorporation of Bodipy-C12-568 was found to be an excellent and sensitive reporter assay to calculate the ACAT activity in living cells.

**Fig. 2.**
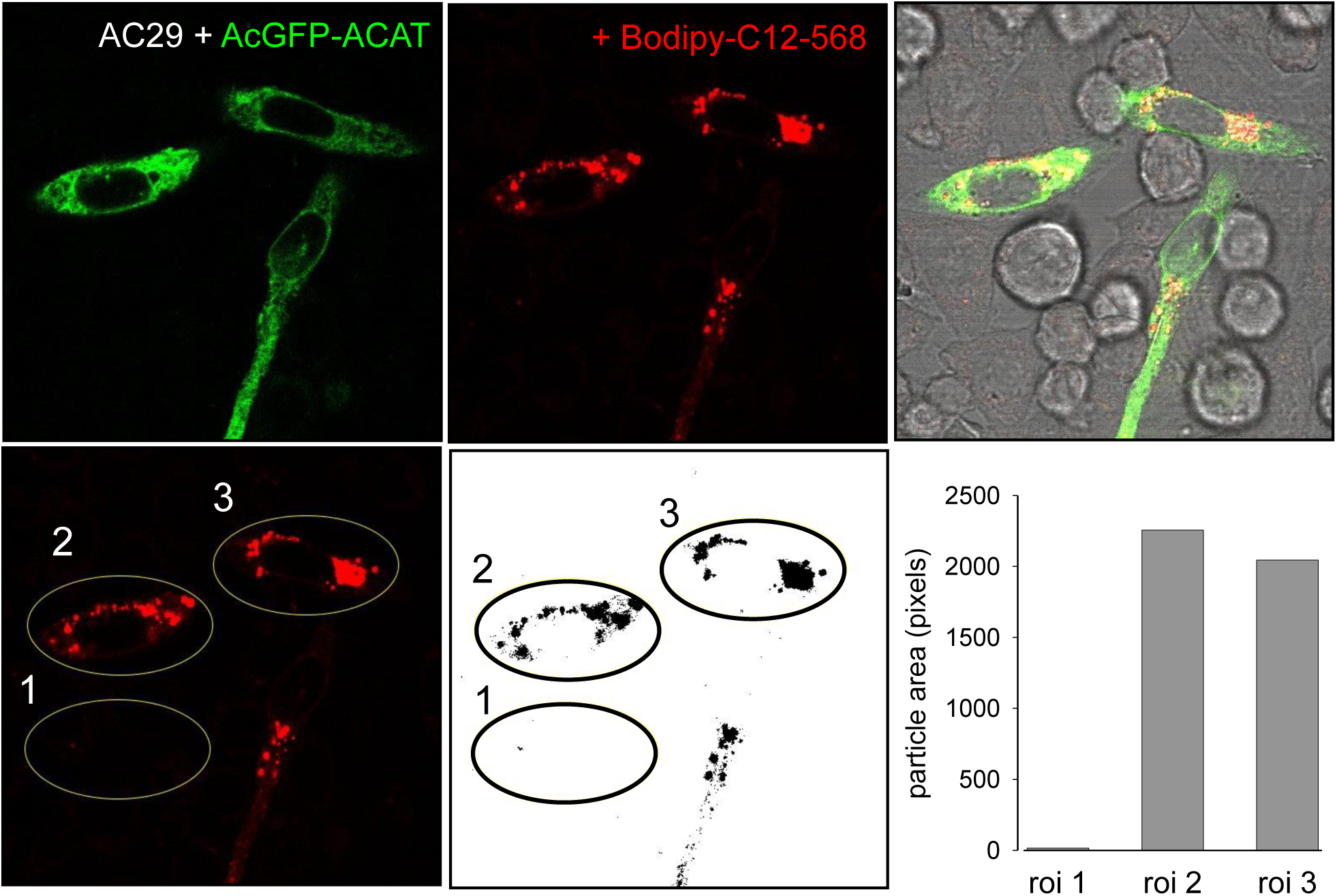
ACAT esterification incorporates fatty acids into lipid droplets. CHO cells that are deficient in ACAT activity (AC29) were transfected with AcGFP-ACAT and were labeled for 10 min with the fatty acid probe Bodipy-C12-568. After a washing step, the cells were incubated in culture medium for 3 h and analyzed by fluorescence microscopy. Upper images: left, AcGFP-ACAT; middle, Bodipy-C12-568; right, overlay including bright field. Lower images: quantification of the Bodipy-C12-568 image using 3 regions of interests (roi); the image was thresholded, binarized and processed for particle counting (Image J, NIH).

### ACAT translocates from the ER to vesicular structures in response to ACAT inhibitors

Under normal physiological conditions, ACAT1 is localized in the widespread ER network. To visualize the ER, the large and flat-shaped fibroblast-like COS-7 cells were stably transfected with EGFP-tagged ACAT1 and were labeled with DChol-MβCD. After several hours of incubation, abundant DChol-containing lipid droplets developed, mostly associated with ER tubules in which ACAT was present (Fig. S1). The majority of the DChol-labeled droplets remain closely connected with the ER as shown in Fig. S1 for one single lipid droplet marked with arrowhead. Formation of lipid droplets most likely occurs through a budding process at the ER and often continue to adhere to the ER membrane after biogenesis [5,42,45,58,63].

To our great surprise, the distribution profile of ACAT changed dramatically after an ACAT inhibitor (iACAT) was added to the cells (Fig. 3). In our first experiments, we have added the ACAT inhibitor SAN 58035 at 0.4 µg/ml for 1 h to HEK-ACAT cells. In nearly all cells, ACAT translocated from the ER (Fig. 3A) to predominantly vesicular structures (Fig. 3B). When iACAT was applied for several hours, ACAT was occasionally observed in cluster-like structures in a couple of cells (Fig. 3C). To exclude the possibility that the ACAT inhibitor induces changes in the ER network in any way, the effect of iACAT was tested on HEK293 cells cotransfected with EGFP-ACAT1 and the ER marker DsRed-ER. While ACAT was again found in vesicular structures, no significant alterations in the ER network were detected (Fig. 3D). In addition, an antibody against the ER resident protein calnexin did not indicate structural changes in the ER in response to ACAT inhibition (Fig. 3E).

**Fig. 3.**
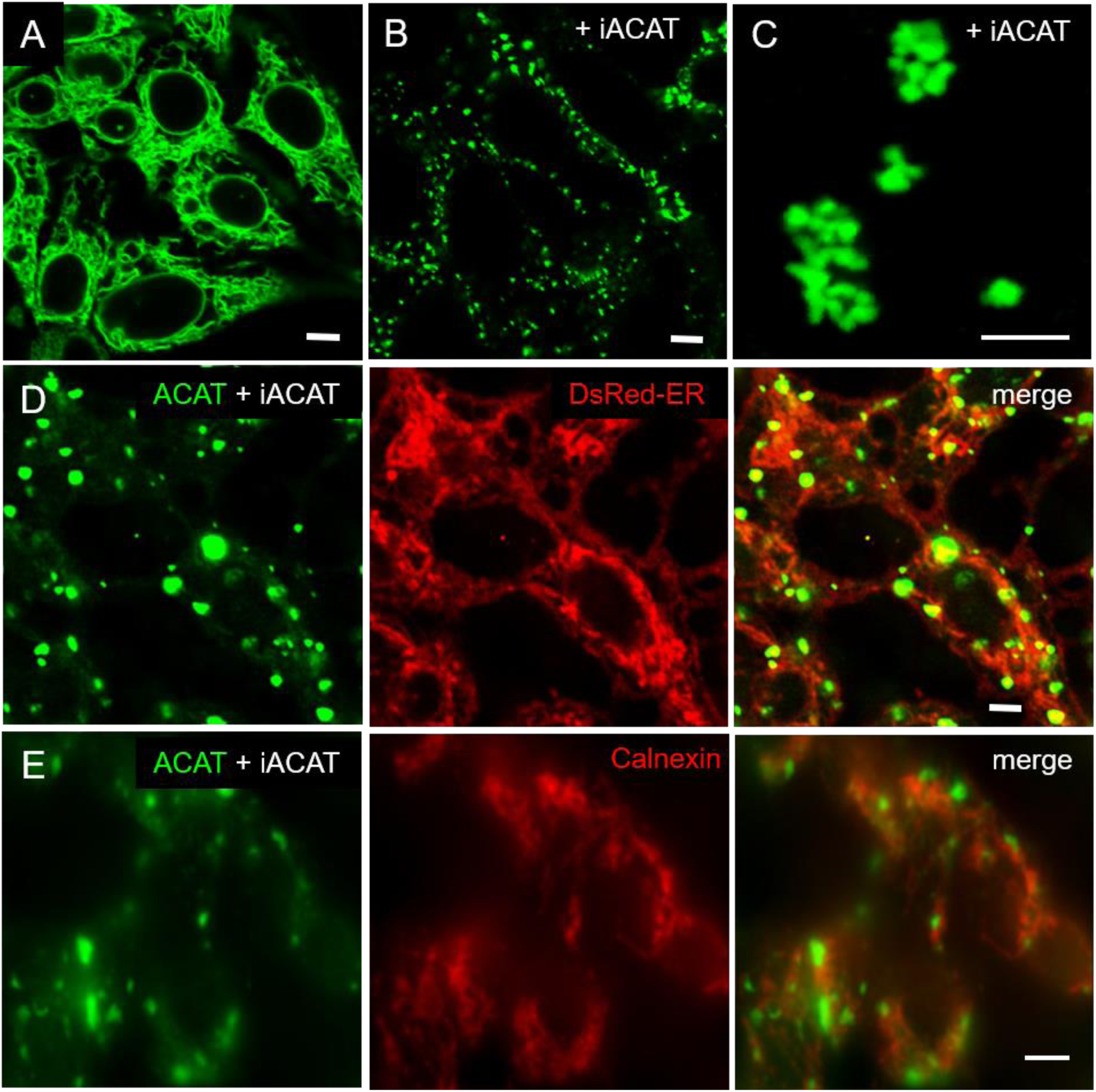
Inhibition of ACAT induces significant changes in its intracellular distribution. HEK-ACAT cells (A-C, E) or HEK cells stably cotransfected with ACATGFP and DsRed-ER (D) were treated (B-E) or not treated (A) for 1 h (B, D, E) or 4 h (C) with the ACAT inhibitor SAN 58035 (0.4 µg/ml) in a microscope chamber at 30°C. Following application of the inhibitor, a predominant vesicular pattern of ACAT (B, D, E) is observed in most cells within minutes. After prolonging the incubation time to several hours, additional cluster-like structures are occasionally found in some cells (C). (E) Formaldehyde-fixed and permeabilized cells were stained with anti-calnexin / anti-mouse-IgG-Cy3. The ER structure (in D and E) did not change significantly after application of iACAT (bars, 5 µm).

First, we assessed whether the translocation of ACAT also occurs in other cells. Both, the CHO-derived AC29 as well as HeLa cells, each transfected with EGFP-tagged ACAT showed a prominent ACAT translocation following the application of ACAT inhibitor SAN 58035 (0.4 µg/ml) for 1 h (data not shown). To rule out the possibility, that adding the EGFP-tag could have led to undesired dimerization/oligomerization of ACAT, we substituted the EGFP-tag by monomeric AcGFP and expressed this construct in AC29 cells. But, AcGFP-tagged ACAT behaved the same way. Following ACAT inhibition, it moved from the ER to vesicles (not shown).

Next, we tested whether the observed translocation behavior of ACAT is a specific response to SAN 58-035 or can also be induced by other ACAT inhibitors. The following additional compounds were used: eflucimibe (=F12511), avasimibe, auraptene, piperine, and tamoxifen. As shown in Table 1, all these compounds were able to cause ACAT translocation, albeit to varying degrees. While SAN 58035, eflucimibe, tamoxifen, and avasimibe were most effective, auraptene and piperine induced ACAT translocation in only a fraction of cells. Among all compounds tested so far, eflucimibe was effective at the lowest concentration and caused a maximum effect in the shortest time. For this reason, we used eflucimibe in most further experiments.

**Table 1.** Effects of ACAT inhibitors on ACAT translocation event. AC29-ACAT cells were treated with the indicated compounds and were then inspected for ACAT1 translocation. The extent of translocation is indicated by symbols: (+++), massive formation of vesicles in virtually every cell; (++), vesicle formation in >80% of cells; (+) weak translocation or translocation in only a fraction of cells; (-) no effect.

| Compound | Treatment (concentration, time) | Translocation |
| --- | --- | --- |
| SAN 58035 | 0.8 μM, 1 h | +++ |
| Eflucimibe | 0.4 μM, 30 min | +++ |
| Tamoxifen | 10 μM, 1 h | +++ |
| Avasimibe | 10 μM, 1 h | ++ |
| Auraptene | 10 μM, 1 h | + |
| Piperine | 20 μM, 1 h | + |

We did not find significant colocalization of the ACAT containing vesicles (or cluster structures) with known organelle markers. As shown in the Figures S2 and S3, the ACAT structures are not colocalizing with lysosomes (Fig. S2A, lysotracker), Golgi (Fig. S2B, anti-Golgin; Fig. S2C, WGA-TR), or mitochondria (Fig. S3A, mitotracker CMTRos). Furthermore, the ACAT vesicles are no ER exit sites as essentially no colocalization with anti-Sec31 immunostaining was observed (Fig. S3C). The highest although still very moderate colocalization was obtained for peroxisomes (with a Pearsońs coefficient of 0.425) employing DsRed-Peroxi as the organelle marker (Fig. S3B).

### ACAT translocation is fast and reversible, but can be prevented by cycloheximide

We now studied the kinetics of ACAT translocation in HEK-ACAT cells in response to eflucimibe. Approximately 5 minutes after ACAT inhibitor administration, both the number and size of ACAT-containing vesicles began to increase (Fig. 4A). About 15 min later, the number of vesicles reached its maximum (Fig. 4A, filled circles). However, the size of the vesicles/clusters further increased for a few more minutes (Fig. 4A, open circles). When eflucimibe was washed off, the number of vesicles began to decrease, albeit at a very low rate. Even after 7 hours of incubation without eflucimibe, nearly 80% of the ACAT vesicles were still present (Fig. 4B, open circles).

**Fig. 4.**
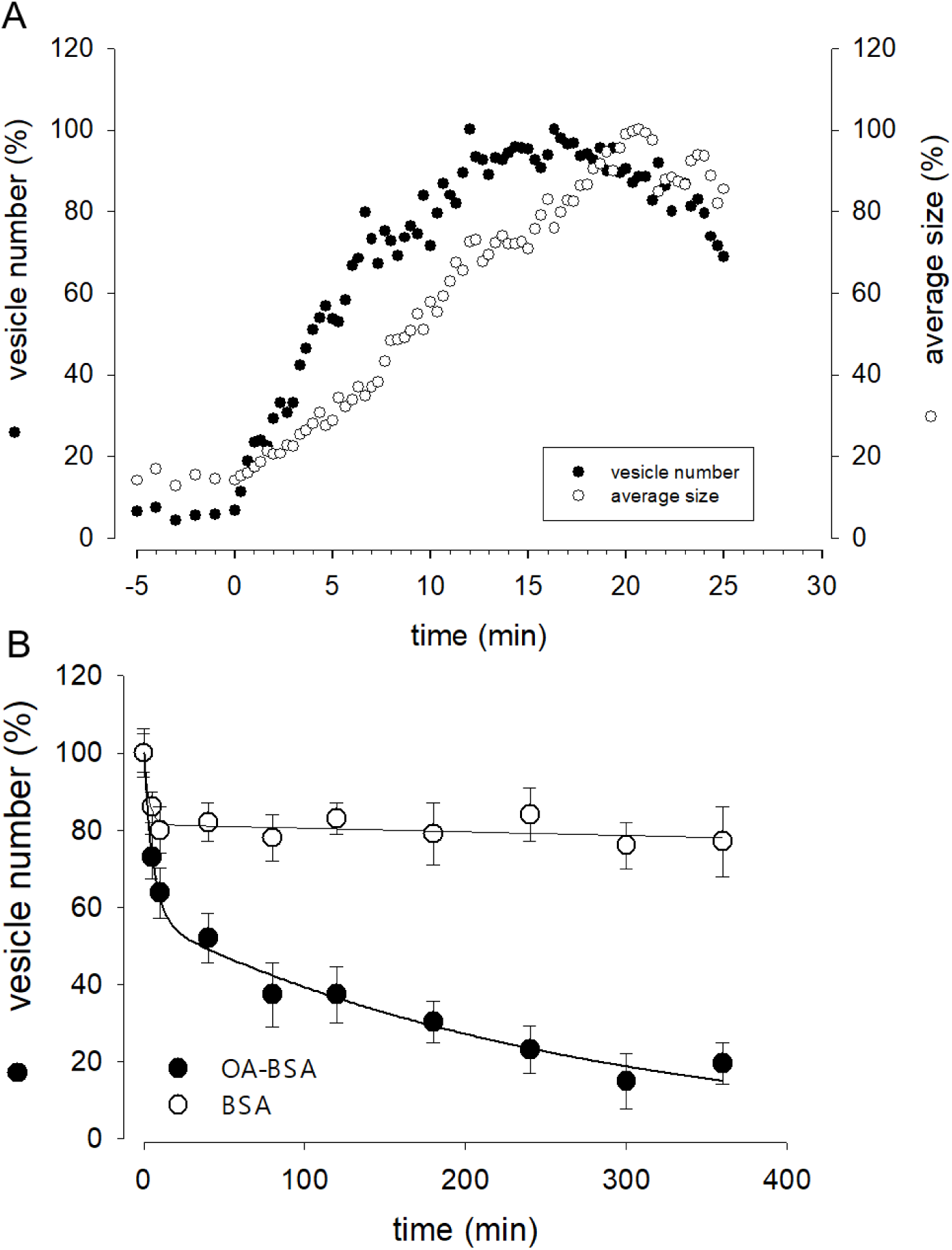
Kinetic and reversibility of the translocation of ACAT. (A) HEK-ACAT cells were cultivated in a temperature controlled microchamber on the microscope stage. The translocation of ACAT was initiated with ACAT inhibitor eflucimibe (0.5 µM). About 5 min after application of eflucimibe, the number (•) and size (ο) of ACAT containing vesicles begin to increase. Images were captured each 20 sec starting at time 0. The vesiculation process was quantified by using the particle counting module of Image J (NIH). (B) Eflucimibe was washed-off after 30 min, and the reversibility of the vesiculation was initiated by addition of BSA (10%) (ο) or BSA-bound oleic acid (400 µM OA in 10% BSA) (•). The number of vesicles decreased as determined by imaging and particle counting at different times. The data were fitted by exponential functions: double exponential (with 4 parameters) for BSA: a = 18.63, b = 0.321, c = 81.457, d = 0.0001184; for OA-BSA: a = 42.83, b = 0.171, c = 56.94, d = 0.0037.

This prompted us to test if certain substances are able to accelerate the reversibility of the translocation process. In the process, we also tried oleic acid as a candidate, as its activated form, oleoyl-CoA, is the cellular substrate of ACAT. Surprisingly, we found that the administration of oleic acid complexed with BSA dramatically increased the re-translocation process of ACAT from vesicular structures into the ER compartment. After less than 5 hours, roughly 80% of the vesicles had disappeared (Fig. 4B, filled circles). In cells that did not receive eflucimibe prior to administration of oleic acid-BSA (for 5 h), ACAT resided unaltered within the reticular ER (not shown). Thus, the translocation process of ACAT was relatively fast, occurring within minutes, and was reversible. In addition, it could be triggered twice in succession on the same cells.

In further experiments, we surprisingly found that the pretreatment of cells with the protein synthesis inhibitor cycloheximide prevented the translocation of ACAT in vesicles. Interestingly, it has been reported that cycloheximide can induce the formation of cholesterol ester-rich lipid droplets [55]. This suggests that ACAT could be somehow activated by cycloheximide. In fact, when HeLa cells were incubated with cycloheximide (for 12 h at 20 µM), the number of lipid droplets significantly increased as shown by the fluorescence marker Bodipy 493/503 (Fig. 5A). After treatment with cycloheximide (CHX), lipid droplets were observed in approximately twice as many cells, with droplet-rich cells (>10 droplets) being as much as 4 times more abundant than in untreated cultures (Fig. 5B). Subsequently, it was verified whether the higher number of lipid droplets in CHX-treated cells was also associated with an increased rate of cholesterol esterification. The lipid extracts of HeLa cells that were incubated with the ACAT substrate DChol in the presence or absence of cycloheximide were analyzed by TLC. Free and esterified DChol was quantitated by fluorimetry. The quantitation of a representative TLC (Fig. 5C, left panel) is shown in the middle panel of the graph. The right panel in Fig. 5C shows the ratio of DChol-Ester to DChol obtained in three experiments with HeLa cells in the presence and absence of cycloheximide, respectively. Significantly higher amounts of DChol esters were formed in the cycloheximide-treated cells than in untreated cells. Finally, the lipid extracts from HeLa cells treated or untreated with cycloheximide were separated by TLC, and the resolved lipid species were visualized by heating after spraying with sulfuric acid:methanol (Fig. 5D). Each of the bands corresponding to free cholesterol (FC), cholesterol esters (CE), and triacylglycerols (TAG) were quantitated and are displayed as mean values ± SD in Fig. 5D (right panel). These data show that, in addition to cholesterol esters, particularly the formation of triacylglycerols was increased in cycloheximide-treated cells, approximately fourfold compared with control cells.

**Fig. 5.**
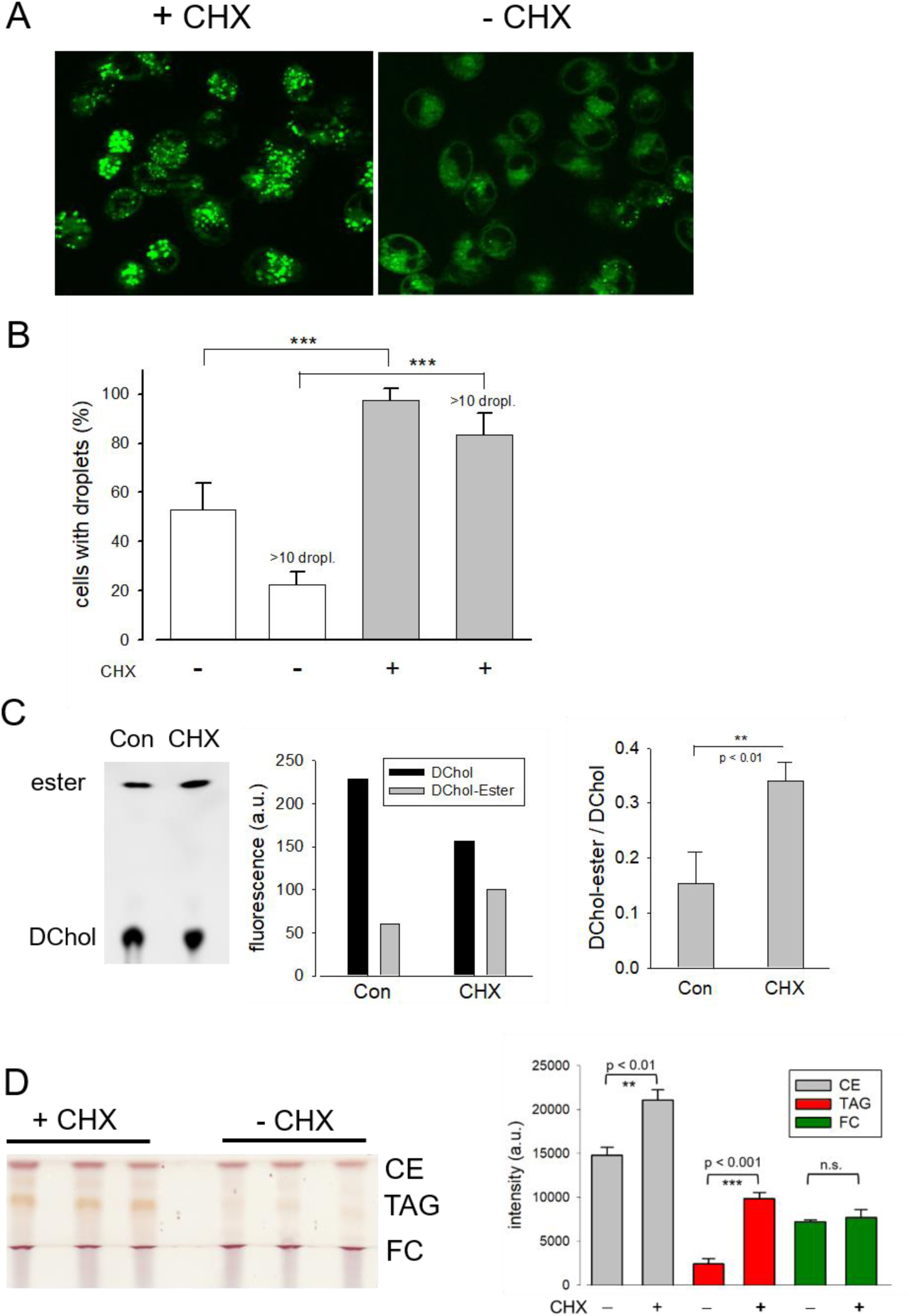
Effect of cycloheximide on lipid droplet formation and cholesterol esterification. (A) HeLa cells grown on glass-bottom dishes were incubated for 12 h in the presence (+ CHX) or absence (-CHX) of cycloheximide (20 µM) in DMEM without FCS. Thereafter, Bodipy 493/503 (4 µM) was added for 30 min. After a washing step, the cells were imaged and droplets were counted (B) as described (n = 80-100 cells; ***, p < 0.001). (C) HeLa cells cultivated in Petri dishes were incubated for 14 h with DChol-MβCD (0.3 mM) in the absence (Con) of presence of cycloheximide (CHX) (20 µM) in DMEM (with FCS). The cells were harvested, and the corresponding lipid extracts were analyzed by TLC. Free and esterified DChol was quantitated by fluorimetry using the imaging device UVP ChemStudio (Analytic-Jena). The quantitation of a representative TLC (left panel) is shown in the middle panel of the graph. The right panel shows the ratio of DChol-Ester to DChol (means ± SD) obtained in three independent experiments. (D) HeLa cells were cultivated in Petri dishes in the absence (Con) or presence of cycloheximide (CHX) (20 µM) in DMEM (with FCS). The cells were harvested, and the corresponding lipid extracts were separated by TLC. The air dried plate was sprayed with sulfuric acid:methanol (1:1 vol%) and heated at 120°C for 5 min. Free cholesterol (FC), cholesteryl ester (CE), and triacylglycerols (TAG) were quantitated by Image J. The data are mean values ± SD (right panel) from the corresponding lipid bands shown in the left panel.

### ACAT translocation can also be induced by other substances and stimuli

Next, it was explored whether other substances or treatments could also induce intracellular translocation events of ACAT. The results are shown in summarized form in Table 2. First, the cholesterol content of HEK-ACAT cells was decreased or increased by incubation with the cholesterol acceptor methyl-β-cyclodextrin (MβCD) or the water-soluble inclusion complex cholesterol-MβCD, respectively. In both cases, a massive translocation of ACAT from ER into vesicles occurred. Since MβCD itself was able to induce ACAT translocation, the specificity of the effect of cholesterol-MβCD was not possible to judge. Therefore, we additionally used cholesterol dissolved in ethanol, which caused a significant ACAT translocation at concentrations >30 µM. A similarly strong effect was achieved when the sterol analogue lanosterol was used. In contrast, the steroid cholesten-3-one was ineffective under the same experimental conditions. Among other substances that are known to affect the cholesterol homeostasis (25-hydroxycholesterol, lovastatin, U18666A) only 25-hydroxycholesterol was able to induce a slight translocation of ACAT. Osmotic stress induced by administration of sucrose also lead to a substantial translocation of ACAT into vesicles. Other substances that are known to interfere with microtubule polymerization (nodocazole), ER to Golgi transport (brefeldin A), or cellular signaling (forskolin, H89, phorbol ester, lysophosphatidic acid, Y27632, wortmannin, U73122) did not alter the intracellular localization of ACAT (Table 2). Interestingly, we also noticed that ACAT translocation was facilitated or promoted when the cells were exposed to a lower temperature (e.g., by transfer from 37°C to room temperature) or cultivated at maximum density.

**Table 2.** Effects of substances or treatments on ACAT translocation. HEK-ACAT cells were treated with the indicated compounds and were then inspected for ACAT1 translocation. The symbols indicate: (++), vesicle formation in >80% of cells; (+) weak translocation or translocation in only a fraction of cells; (-) no effect.

| Compound | Effect | Treatment | Translocation |
| --- | --- | --- | --- |
| Cholesterol-MβCD | cholesterol increase | 0.3 mM, 30 min | ++ |
| Methyl-β-cyclodextrin (MβCD) | cholesterol decrease | 20 mM, 10 min | ++ |
| Cholesterol (EtOH) | cholesterol increase | 30 μM, 2 h | + |
| Lanosterol (EtOH) | lanosterol increase | 30 μM, 2 h | + |
| 4-Cholestene-3-one (EtOH) | 4-cholestene-3-one increase | 30 μM, 2 h | - |
| 25-Hydroxycholesterol | cholesterol activation | 20 μM, 4 h | + |
| Lovastatin | cholesterol synthesis inhibitor | 5 μM, 4 h | - |
| U18666A | cholesterol trafficking inhibitor | 5 μM, 4 h | - |
| Nodocazole | blocks microtubuli polymerization | 1 μM, 4 h | - |
| Brefeldin A | ER to Golgi transport inhibition | 20 μM, 4 h | - |
| Forskolin | adenylyl cyclase activation | 100 μM, 1 h | - |
| H89 | protein kinase A inhibition | 20 μM, 1 h | - |
| Phorbol 12-myristate 13-acetate | protein kinase C activation | 100 μM, 1 h | - |
| Lysophosphatidic acid | rho kinase activation | 10 μM, 30 min | - |
| Y27632 | rho kinase inhibition | 100 μM, 1 h | - |
| Wortmannin | phosphoinositide 3-kinase inhibition | 100 μM, 1 h | - |
| U73122 | phospholipase C inhibition | 100 μM, 1 h | - |
| Sucrose | osmotic stress | 0.45 M, 10 min | ++ |

### The translocation process is not accompanied by changes in electrophoretic mobility of ACAT

The question arises whether the translocation of ACAT is accompanied by molecular alterations of the protein such as oligomerization, aggregation or proteolysis. First, the migration behavior of endogenous ACAT in SDS-PAGE (Fig. 6A, left lanes) was analyzed. HEK293 cells treated or not treated with eflucimibe (0.5 µM, 1 h) were homogenized, and the proteins were separated by SDS-PAGE, followed by immunoblotting with anti-ACAT/SOAT antibody. The endogenous ∼50 kDa monomeric anti-ACAT species is running at a slightly lower than predicted molecular weight (∼64 kDa). This differential migration pattern of ACAT is known from earlier studies and was assumed to be caused by the high IEP (∼8.7) of the protein [13]. In the presence of eflucimibe, the electrophoretic mobility of endogenous ACAT did not change (Fig. 6A, left lanes). Even when protein analysis was performed using native/non-denaturing conditions with HEK-ACAT cells, eflucimibe induced no alterations in the gel mobility of ACAT (Fig. 6A, right lanes).

**Fig. 6.**
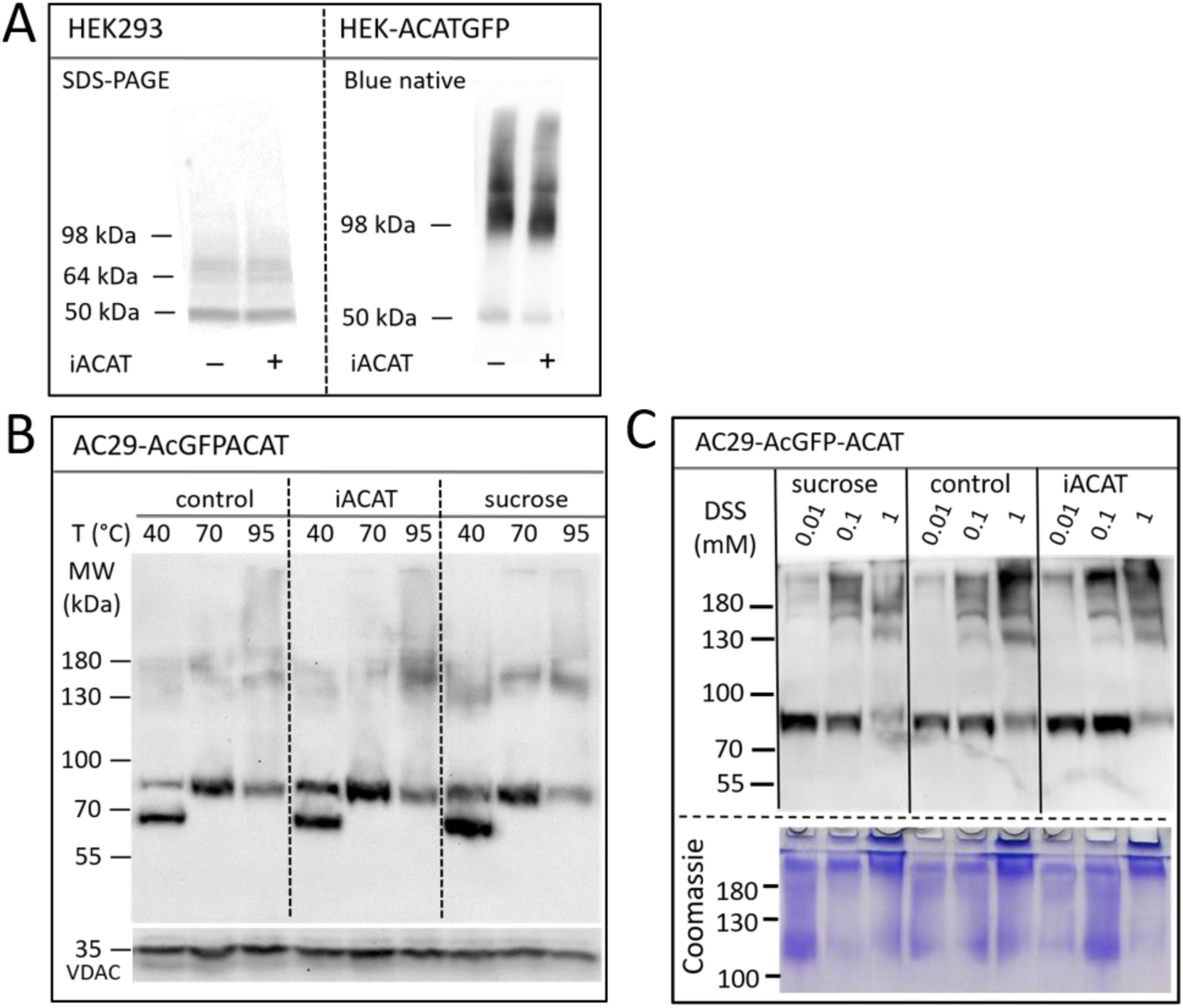
Analysis of the electrophoretic mobility of ACAT in dependence of the translocation inducers eflucimibe (iACAT) and/or sucrose. (A) HEK293 cells transfected or not transfected with ACATGFP were incubated for 1 h in the presence or absence of eflucimibe (0.5 µM). Membranes of these cells were prepared and the proteins were separated by denaturing SDS-PAGE or by non-denaturing Blue native gel electrophoresis, followed by immunoblotting. (B) AC29-ACAT cells were incubated in the absence (control) or presence of sucrose or eflucimibe (iACAT) for 1 h. The cells were homogenized and the membrane proteins were analyzed by SDS-PAGE and immunoblotting. Prior to gel electrophoresis, samples were heated for 10 min at either 40°C, 70°C or 95°C in SDS loading buffer. Immunodetection of ACAT was performed using anti-ACAT/SOAT antibody, while VDAC (∼35 kDa) was used as a loading control. (C) AC29-ACAT cells were treated or not treated (control) with sucrose or eflucimibe (iACAT) for 1 h. Then, the membrane-permeable aminoreactive crosslinker disuccinimidyl suberate (DSS) was added to the cells at increasing concentrations (0.01 mM, 0.1 mM, 1 mM) for 30 min at 37°C. After washing-off the crosslinker, the cells were homogenized and the membrane proteins were analyzed by SDS-PAGE and immunoblotting using anti-ACAT/SOAT antibody.

We also studied the electrophoretic mobility of ACAT using AC29-ACAT cells and incubated in the absence (control) or presence of sucrose (450 mM) or eflucimibe (iACAT) (0.5 µM, 1 h). The membrane proteins were analyzed by SDS-PAGE and immunoblotting (Fig. 6B). The migration pattern of ACAT in SDS-PAGE was markedly affected by the temperature at which the samples were incubated prior to gel electrophoresis. Heating the samples at 40°C, 70°C or 95°C for 10 min in SDS loading buffer revealed substantial differences in their electrophoretic mobility (Fig. 6B). Monomeric AcGFP-tagged ACAT was running at a ∼85 kDa species which was again slightly lower than its calculated molecular weight (∼94 kDa). In addition to this species, a lower molecular ∼70 kDa form appeared at 40°C and disappeared at higher temperatures (70°C and higher). At 95°C, the intensity of the monomeric ∼85 kDa ACAT species was decreasing while at the same time higher molecular ACAT dimers and oligomers appeared. However, when compared with untreated cells, neither eflucimibe nor sucrose led to substantial alterations in the gel-mobility of ACAT.

Finally, the electrophoretic mobility of ACAT was explored under crosslinking conditions. AC29-ACAT cells were incubated in the absence or presence of sucrose (450 mM) or eflucimibe (iACAT) (0.5 µM) for 1 h. Then, the membrane-permeable aminoreactive crosslinker disuccinimidyl suberate (DSS) was added to the cells at increasing concentrations (0.01 mM, 0.1 mM, 1 mM) for 30 min at 37°C. The cells were washed with PBS, homogenized by sonication, and centrifuged by 30,000g for 30 min (4°C). The pelleted proteins were dissolved in SDS loading buffer and were treated for 10 min at 70°C prior to gel loading, separation by SDS-PAGE, and immunodetection using anti-ACAT/SOAT antibody. To demonstrate the successful crosslinking with DSS, the remaining proteins of the SDS gel were stained by Coomassie. As shown in Fig. 6C (lower panel), DSS at concentrations of 1 mM produced high molecular protein aggregates that partly remained in the stacking gel. In the presence of increasing concentrations of the crosslinker DSS, the intensity of the monomeric anti-ACAT species was decreasing while at the same time higher molecular anti-ACAT species (dimers and oligomers) appeared as expected (Fig. 6C). However, even after chemical crosslinking, ACAT translocation was not associated with changes in the electrophoretic mobility of the protein. In addition, no signs of proteolysis during the translocation process of ACAT were observed.

### Translocated ACAT can esterify cholesterol efficiently

Is ACAT associated with a change or loss of its enzymatic activity in its vesicular environment? To address this question, we used the fluorescent cholesterol reporter DChol and established an *in vitro* assay to measure the activity of ACAT. When both substrates, DChol (10 µM) and oleoyl-CoA (0.8 - 64 µM) were added to microsomes from AC29-ACAT cells, an esterification of DChol was clearly observed (Fig. 7A). Increasing the substrate concentration of oleoyl-CoA resulted in increasing rates of DChol esterification. At high concentrations of oleoyl-CoA, saturation of the enzyme was observed as expected (Fig. 7B). Using this assay, the activity of vesicular *versus* non-vesicular ACAT was compared. Of course, an ACAT inhibitor as the stimulus for ACAT translocation could not be employed in these experiments. For this purpose, hyperosmotic sucrose treatment of cells was used as the stimulus to induce the translocation of ACAT. The experiments were performed in AC29-ACAT cells as well as in HeLa cells endogenously expressing ACAT. The cells were incubated for 1 h in the absence (control) or presence of sucrose (450 mM in DMEM medium). The ACAT translocation in sucrose treated cells was confirmed by microscopical inspection. All cells were harvested by a cell scraper and their microsomes were prepared. The ACAT activities of these microsomes were measured by the *in vitro* assay mentioned above. Following lipid extraction and separation by TLC, the DChol and DChol-oleate bands were quantified by fluorimetry. The band intensities of DChol, DChol-ester as well as the ratios of DChol-ester to DChol were calculated (Table 3). The data are graphically displayed in Fig. 8, with the control cell data set to 100%. In microsomes from HeLa cells, nearly 5-fold lower DChol-ester amounts were found as compared with microsomes from AC29-ACAT cells (Table 3). This is certainly due to the higher expression level in AC29-ACAT cells. More importantly, it was found that the ACAT activities in microsomes from both sucrose treated cells were at least as high as those obtained in microsomes from untreated cells. From this, it can concluded that ACAT, which is translocated into vesicles in response to osmotic stress, is fully activatable when sufficient substrate is available.

**Fig. 7.**
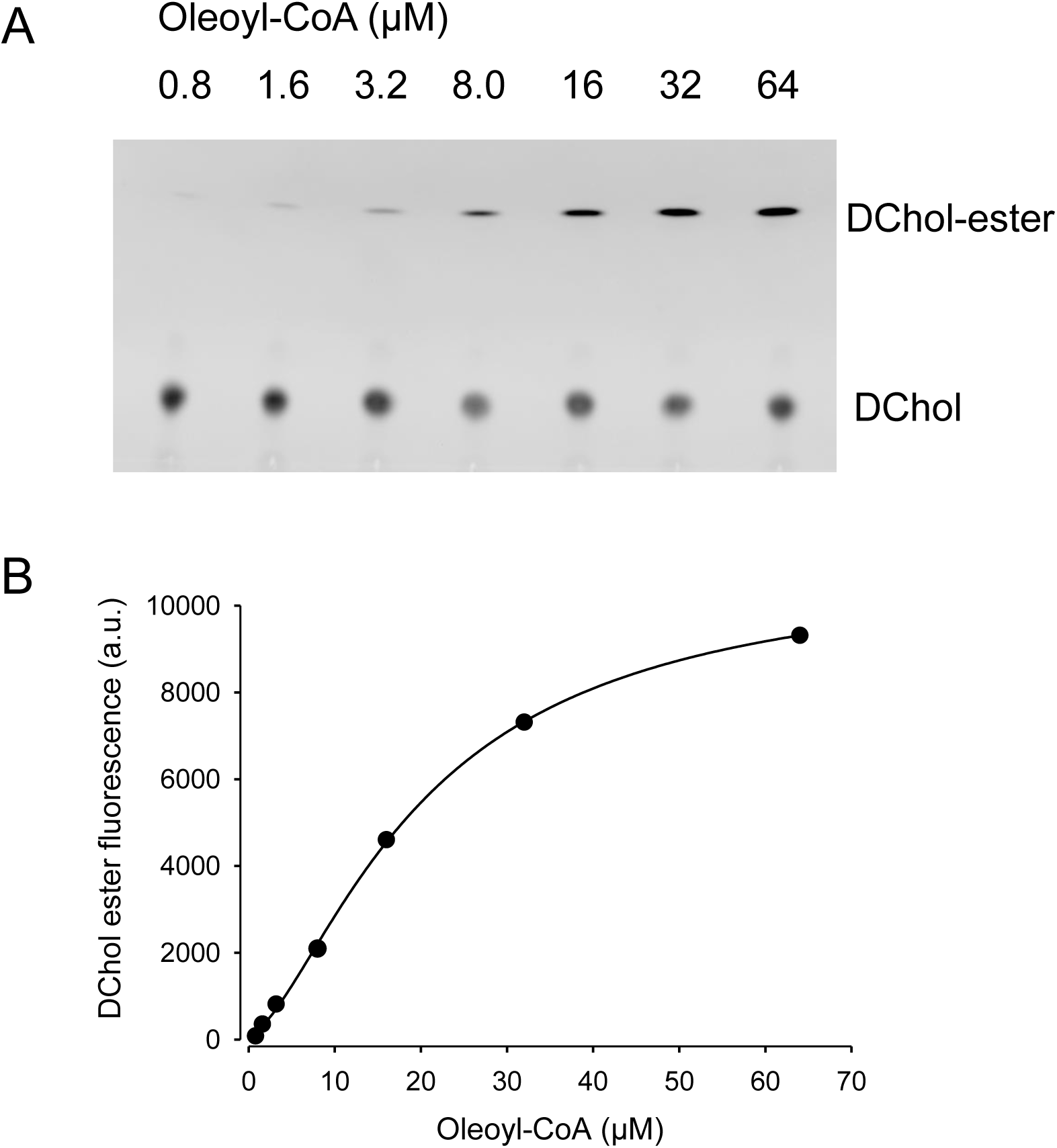
ACAT activity assay in microsomes. (A) Microsomes prepared from AC29-ACAT cells (2 x 10^6^ cells) were resuspended in 300 µl assay buffer (20 mM Hepes, pH 6.8, 2 M KCl). After addition of 1 µl DChol-MβCD (3 mM stock), the assay was mixed and incubated for 5 min at 25°C. The esterification reaction was initiated by addition of 1 µl oleoyl-CoA (final 0.8 – 64 µM from increasing stock solutions). After incubation for 2 h at 37°C, the lipids were extracted and separated by TLC on silica gel G plates using petrolether:diethylether:acetic acid (8:2:1, v/v/v) as the solvent system. (B) The DChol ester bands were quantified by fluorimetry. The data were best fit by a 4-parameter logistic function yielding an EC_50_ of 20.1 µM for oleoyl-CoA (other parameters: max 10877, min 97.3, slope 1.5).

**Fig. 8.**
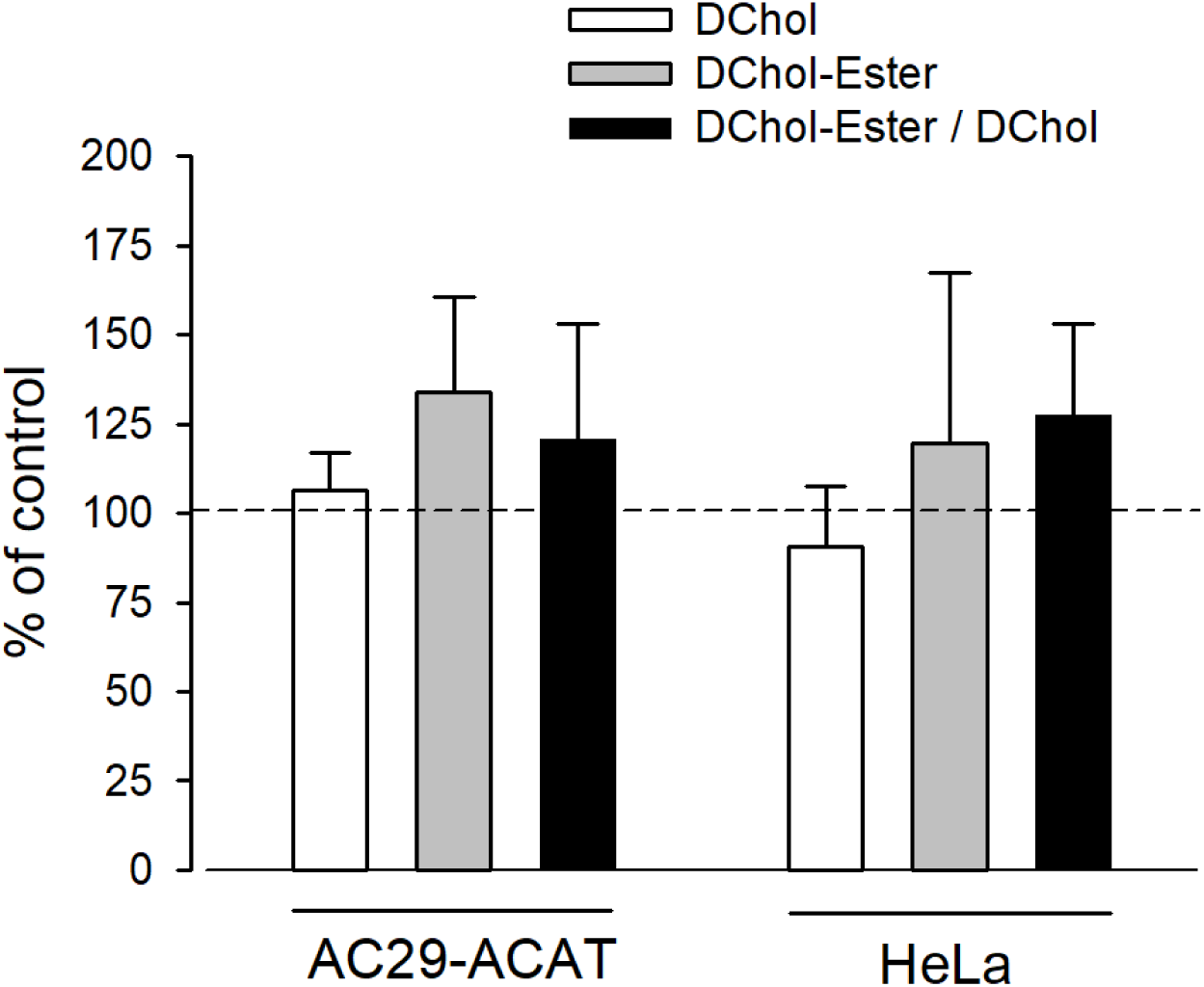
Analysis of sucrose-induced ACAT translocation on the activity of ACAT. AC29-ACAT and HeLa cells were cultivated in 6 cm dishes and were incubated in the absence (control) of presence of sucrose (450 mM) for 1 h at 37°C. The cells were harvested, lysed by sonication and subjected to microsomal preparation. The microsomes were resuspended in 300 µl assay buffer (20 mM Hepes, pH 6.8, 2 M KCl). After addition of 1 µl DChol-MβCD (final 10 µM), the assay was mixed and incubated for 5 min at 25°C. The esterification reaction was initiated by addition of 1 µl oleoyl-CoA (final 20 µM). After incubation for 2 h at 37°C the lipids were extracted and separated by TLC. DChol and DChol ester bands were quantified by fluorimetry. The band intensities of DChol (white bars), DChol-ester (grey bars) as well as the ratios of DChol-ester to DChol (black bars) were calculated. The data of the untreated cells were set to 100%. The respective values of the sucrose-treated cells are graphically shown as a percentage of this (± SD, n=3) (data in Table 3). The differences between control and sucrose treatment remained below the significance level in case of both DChol and DChol-ester.

**Table 3.** ACAT activity in microsomes from AC29-ACAT and HeLa cells treated or not treated with sucrose (450 mM for 1 h). The cells were harvested, lysed by sonication and microsomes were prepared as described. The ACAT activity in these microsomes was measured using the substrates DChol and oleoyl-CoA. After extraction and separation of the lipids by TLC, the DChol and DChol ester bands were quantified by fluorimetry. The fluorescence intensities (a.u.) are given as means ± SD obtained from three experiments.

| Cell | Parameter | Control | Sucrose |
| --- | --- | --- | --- |
| AC29-ACAT | DChol | 19994 ± 1344 | 22703 ± 1477 |
| | DChol-ester | 3523 ± 1176 | 4702 ± 1622 |
| | DChol-ester / DChol | 0.177 ± 0.058 | 0.205 ± 0.055 |
| HeLa | DChol | 21756 ± 5125 | 19115 ± 1113 |
| | DChol-ester | 951 ± 505 | 1027 ± 340 |
| | DChol-ester / DChol | 0.042 ± 0.01 | 0.053 ± 0.015 |

## DISCUSSION

This study was initiated by the incidental observation that commonly used ACAT inhibitors induce a fast (within minutes) intracellular translocation of ACAT from the extensive network of the endoplasmic reticulum into a vesicular compartment. This occurred in all cells studied so far. In the process, we also discovered that a number of other substances or treatments can induce such a ACAT translocation, e.g., cholesterol or lanosterol at concentrations >30 µM, 25-hydroxycholesterol, and certain stressors such as osmotic stress induced by sucrose, exposition of the cells to lower temperatures or cultivation of the cells at maximum density. While the majority of ACAT vesicles in the present study showed no clear assignment to a defined compartment, a small population of these vesicles was located in close proximity to peroxisomes. This is of potential interest because membrane contacts between peroxisomes and lysosomes represent an important transport pathway for LDL-derived cholesterol. Disruption of these contacts induces massive cholesterol accumulation in lysosomes, as observed in fibroblasts from patients with different types of peroxisomal disorders [15]. Peroxisomes play a central role in the cellular lipid metabolism. For example, the β-oxidation of fatty acids, the pre-squalene segment of the cholesterol biosynthesis and the formation of bile acids take place in this organelle [12]. Principally, at such peroxisomal contact sites, ACAT could have access to LDL-derived cholesterol and to its co-substrate acyl-CoA, which can possibly be provided by acyl-CoA synthetases localized to the surface of peroxisomes [14]. However, this issue requires further investigation.

Of particular importance is our finding that translocation of ACAT from the extensive ER network into vesicular structures is a reversible process that under certain conditions can even be triggered twice in succession on cells, for example, after application and removal of the ACAT inhibitor eflucimibe. Remarkably, oleic acid was able to greatly accelerate the re-translocation of ACAT from the vesicles back to the ER. Considering that oleic acid in its activated form (oleoyl-CoA) is a preferred fatty acid substrate of ACAT, it could be speculated that this translocation process might be of physiological relevance. Treatment of cells with oleic acid is commonly used to induce the formation of lipid droplets, i.e., the storage of lipids. The development of lipid droplets requires ACAT whose activity depends on the availability of both, cholesterol and long-chain fatty acyl-CoA. The latter are produced by acyl-CoA synthetases localised on peroxisomal, ER and mitochondrial outer membranes [21]. The re-translocation of ACAT from vesicles into the ER network in response to oleic acid, i.e., under lipid storage conditions appears reasonable in view of the following aspect. The extensive network of the ER in the cell, in which ACAT is evenly distributed under normal physiological conditions, provides an enormous surface area and ensures that cholesterol can be efficiently used as substrate. ACAT immediately esterifies ‘activ’ cholesterol molecules that flow from the plasma membrane into this giant ER compartment using Aster transfer proteins [43,53]. Subsequently, the esters are deposited into lipid droplets. This process serves to store energy and also counteracts the accumulation of free cholesterol, that is toxic to the cell. With an adequate supply of cholesterol and oleoyl-CoA, the distribution of ACAT in the ER network seems to be appropriate and well adapted. But what could be the benefit of localising ACAT in vesicles? A regulatory function would be obvious. In particular, this way the enzyme would not be exposed to the non-vesicular influx of cholesterol from the plasma membrane, which is a major pathway of cholesterol entry into the cell [28,40]. The availability and accessibility of ACAT to its substrates could be completely different in these vesicular structures than within the ER network. It is possible that the enzyme gains access to other vesicular transport routes within the cell. At this point, it is also worth remembering that ACAT has two different tasks to fulfil in cholesterol metabolism. It must provide cholesterol esters not only for storage in lipid droplets but also for the formation of lipoproteins. How these two so different functions of the enzyme are regulated is still completely unclear. Regulation of the enzymatic activity through a differential localization would be a conceivable scenario. It is also possible that the translocation of ACAT into vesicles is associated with the strengthening of ER-mitochondria connectivity in response to ACAT inhibition as recently reported [25].

Previous studies have already reported localization of ACAT1 in vesicular structures [29,31,52]. In macrophages for example, a fraction of ACAT1 immunoreactivity was present in ER-derived, perinuclear structures near the trans-Golgi network and the endocytic recycling compartment [31]. Interestingly, ACAT1 positive vesicles have also been observed following cholesterol loading [29,52]. In isolated ER-derived vesicles, Sakashita et al. (2010) [52] detected an increased ACAT activity and speculated in this regard on a mechanism by which ACAT might upregulate its enzymatic activity without having to produce more protein. Actually, we also observed here the formation of ACAT containing vesicles in response to cholesterol. Moreover, 25-hydroxycholesterol, which is known to activate the esterification of cholesterol, was able to induce the translocation of ACAT [19,61]. This suggests that vesicular ACAT can be associated with enhanced enzymatic activity. Interestingly, ACAT translocation also spontaneously occurred when cells approached confluency. Cell growth appears to be correlated with the esterification of cholesterol [3,18]. In endothelial cells, both cholesterol and cholesteryl ester increased markedly as cells reached confluency [16]. The same was observed for CHO cells [37,57]. Thus, ACAT activity appears to be upregulated with increasing cell density, again suggesting that the enzyme functions very efficiently in its vesicular environment.

Despite some efforts, we failed to detect the translocation process of ACAT in cells endogenously expressing the enzyme (e.g., in HeLa cells) by immunocytochemistry using the commercially available antibodies, although these antibodies were suitable for the detection of ACAT in immunoblots. In general, we found that in HEK-ACAT cells expressing very low levels of ACAT, translocation of ACAT was less pronounced and involved fewer and smaller vesicles than in cells more highly expressing the enzyme. Thus, it appears that sufficient numbers of ACAT molecules are required to clearly visualize this translocation process microscopically. According to biochemical and structural data, ACAT was proposed to form tetramers [24,46,50,51]. Thus, it is reasonable to speculate that ACAT translocation could be related to dimerization/oligomerization of the protein. Assuming that ACAT dimers/tetramers are in equilibrium with higher-order oligomers in the ER membrane, increased expression of ACAT should also enhance the formation of oligomeric complexes. So, it cannot be excluded that higher-order oligomers of ACAT can somehow contribute to vesicle formation in ACAT overexpressing cells. Although we did detect some higher molecular weight ACAT complexes in native blue gels and in SDS-PAGE after cross-linking, ACAT translocation was not associated with significant changes in protein electrophoretic mobility.

There is some evidence that at least one other unknown protein is required for ACAT translocation since the protein synthesis inhibitor cycloheximide was able to block this translocation process. The cycloheximide effect is interesting because it refers to earlier observations that remained unexplained. ACAT activity was found to be enhanced by cycloheximide treatment which lead to the conclusion that there is a short-lived protein inhibitor of ACAT-mediated cholesterol esterification [7,56]. It was also reported that cycloheximide (and other translation inhibitors) are able to induce the formation of cholesterol ester-rich lipid droplets [55], a finding which could be confirmed here for HeLa cells. The results of our study may provide a simple explanation for this observation, because cycloheximide-induced prevention of ACAT translocation could lead to persistent esterification of plasma membrane cholesterol and thus to an increased lipid droplet formation. As mentioned above, this non-vesicular cholesterol influx into the ER is regarded as the major pathway of cholesterol entry into the cell. The molecular machinery that can be induced by cycloheximide appears to be complex. So, it was recently reported that treatment of HeLa cells with cycloheximide and other translation elongation inhibitors are able to activate the MondoA pathway [67]. As a member of the MLX family of glucose-sensing transcription factors, MondoA is not only a key regulator of the energy metabolism [30] but also plays an important role in the regulation of cholesterol/steroid metabolism [41,65].

ACAT enzymes are crucial components of cholesterol metabolism, and inhibition of their activity has great therapeutic potential, e.g., against hyperlipidemia, atherosclerosis, Alzheimeŕs disease, Niemann Pick C disease, viral activity, and tumor growth [2,11,23,26,27,35,38,48,49,68,70,71]. For example, avasimibe, one of the most widely used ACAT inhibitors and a potent inducer of ACAT translocation in this study, has recently emerged as a promising anti-tumor compound [64,69,70,72]. We show here that ACAT inhibition is associated with rapid changes of the intracellular distribution of ACAT. However, future studies are needed to confirm these observations, particularly in cells endogenously expressing ACAT/SOAT.

## MATERIALS AND METHODS

### Materials

The substances were from the following suppliers: bovine serum albumin (BSA, fraction V), Biomol (Germany); lipofectamine 2000, ThermoFisher; Rotifect Plus, Carl Roth; SAN 58035, Novartis; eflucimibe (= F12511), gift from Institute de Recherche Pierre Fabry (France); tamoxifen, avasimibe, auraptene, piperine, Merck; T4 DNA Polymerase, Phusion High-fidelity DNA polymerase (PCR), DNA ligase, and all restriction enzymes, New England Biolabs (NEB); Mowiol 4-88, calbiochem (Bad Soden, Germany); disuccinimidyl suberate (DSS), Sigma; DChol, synthesized in our group; Mitotracker orange CMTMRos, lysotracker Red, ThermoFisher; wheat germ agglutinin, Texas Red-X Conjugate (WGA-TR), Bodipy 493/503, ThermoFisher; i-block (Tropix, Applied Biosystems). Antibodies: anti-calnexin, Santa Cruz, #23954; anti-ACAT/SOAT, Santa-Cruz, #sc69836; anti-mouse IgGκ-BP, Santa Cruz, #sc516102 (HRP-conjugated); anti-mouse-Cy3, Sigma, #C2181; anti-VDAC, Santa Cruz, #390996; anti-Golgin-97, ThermoFisher, #A21270; anti-sec31A, BD Transduction laboratories, #612350. All other substances, unless otherwise stated, were from Merck (Darmstadt, Germany).

### Preparation of oleic acid-BSA and sterol-MβCD complexes

Oleic acid (OA) (400 µM) was dissolved in ethanol and evaporated under nitrogen to a thin film. Then, 10% BSA in phosphate-buffered saline (PBS) at 45°C was added to the tube to obtain the desired oleic acid-BSA complex. After the fatty acid was completely dissolved, the solution was cooled, filtered, and stored in aliquots at 4°C. Inclusion complexes of sterols and MβCD were performed as described [66]. For preparation of DChol-MβCD, the steroid (final concentration of 3 mM) was added to an aqueous solution of MβCD (30 mM) in a 2 ml tube. The mixture was overlaid with N_2_ and was vortexed for 24 h at 30°C. The solution was centrifuged to remove insoluble material before use.

### Expression plasmids and cloning

The cDNA encoding human ACAT1 was obtained from RZPD (Berlin, Germany) and was amplified by PCR using the primers 1_for (5′-GCATTTGTACAAGCCCGGCGTGGGTGAAGAGAAGATGTCTCTA-3′) and 2_rev (5′-GCATTGCGGCCGCCTAAAACACGTAACGACAAGTCCAG-3′), digested with BsrGI and NotI (∼1.7 kb fragment). To obtain the plasmid pcDNA5/FRT-GFP, pEGFP-N3 (Clontech) was cut with NotI, blunted with T4 Polymerase and further digested with NheI. The resulting ∼ 700 bp fragment was ligated with pcDNA5/FRT cut with NheI + EcoRV. To construct pcDNA5/FRT-GFP-ACAT1, pcDNA5/FRT-GFP restricted with BsrGI and NotI was ligated with the PCR product of hACAT1 that has been digested with BsrG1 + NotI. The plasmid pAcGFP-ACAT1 was a ligation product of the vector pcDNA5/FRT-GFP-ACAT1, digested with BsrGI, blunted with T4 DNA polymerase, cut with NheI, and the insert encoding AcGFP that was isolated from pAc-GFP-Sec61beta (from Tom Rapoport, Addgene #15108) restricted with BglII, blunted with T4 DNA polymerase, and digested with NheI. To create pcDNA-AcGFP-ACAT1, the vector pfmOTRf [22] cut with HindIII, blunted with T4 DNA polymerase, and digested with XhoI was ligated with the AcGFP encoding insert isolated from pAcGFP-ACAT1, cut with NheI, blunted with T4 DNA polymerase, and restricted with XhoI. To obtain pcDNA5/FRT-BTBhACAT1, the sequence encoding GFP was removed and substituted by the BTB motif. The vector pcDNA/FRT-GFPhACAT1 was digested with BsrGI and NheI and was ligated with an adapter cDNA encoding the BTB-motif. The adapter was created by annealing the two oligonucleotides BTB_1 (5′-CTAGCACCATGGGATGGAGATACTACGAGAGCTCCCTGGAGCCCTACCCTGACGGCAT-3′) and BTB_2 (5′-GTACATGCCGTCAGGGTAGGG CTCCAGGGAGCTCTCGTAGTATCTCCATCCCATGGTG-3′). To create the plasmid pDsRed-Peroxi, the calreticulin signal sequence of pDsRed2-ER (Clontech) was removed by cleavage with NheI and BglII. The vector was ligated with the insert obtained by PCR using the template pDsRed2-ER, the oligonucleotides SKL_for (5′-GCCGGCTAGCATGGCCTCCTCCGAGAAC-3′) and SKL_rev (5′-GGCAGATCTACAGCTTGGACTTGTACGATCTCAGGAACAGGTGGTGGCGGCC-3′), followed by digestion with NheI and BglII.

### Cell culture, transfection, and membrane preparation

HEK293, HepG2, COS-7 and HeLa cells were cultivated in DMEM plus 10% FCS. CHO-K1 and AC29 cells were grown in DMEM/Ham’s F12 medium with 10% FCS. Transfections were performed with lipofectamine 2000 or Rotifect Plus according to the instructions of the supplier. To prepare membranes, the cells were harvested by centrifugation at 1000g. Then, the cells were resuspended in hypotonic lysis buffer (20 mM Hepes, 7.4, 5 mM MgCl_2_) and were homogenized by a Potter-Elvehjem Homogenizer or by sonication (Branson sonifier). In some experiments, cell debris and nuclei were removed by low-speed centrifugation (5000 rpm). The membranes were pelleted for 30 min at 31,000g (4°C), and resuspended in lysis buffer or in an appropriate working buffer. Protein was determined by the Bradford assay (Roti-Quant) using bovine serum albumin as standard.

### Lipid extraction and analysis

All lipid extractions were performed by the method of Bligh and Dyer [4], slightly modified as described [39]. To measure the esterification of DChol, the cells were grown on Petri dishes for various times with culture medium / 10% FCS containing 7.5 µM DChol-MβCD. After washing-off the cholesterol donor, the cells were incubated in culture medium at 37°C for different times in the presence or absence of the indicated ACAT inhibitor (e.g., eflucimibe). The cells were harvested by scraping and their lipids were extracted. The lipids were dissolved in chloroform, separated by silica gel thin layer chromatography (TLC) using heptane/diethyl ether/methanol/acidic acid (80:35:3:2, vol %) as mobile phase. The fluorescence was detected under UV and was imaged with a CCD camera (UVP ChemStudio, Analytic-Jena). Quantification of the data were performed by the software VisionWorks or Image J (Fuji version). Cholesterol was assayed spectrophotometrically using R-Biopharm kit (Darmstadt, Germany) performed in a microscale dimension.

### Fluorescence microscopy and imaging

In most experiments, cells grown on Petri dishes were transferred to 3 cm glass-bottom dishes (MatTek, U.S.A.) for imaging. In some experiments, cells were grown on glass coverslips in Petri dishes. These coverslips were placed in an imaging chamber (Life Imaging Services, Switzerland), which was inserted into an adapter on the microscope stage and temperature controlled with a Peltier device. Cells were imaged using a Zeiss Axiovert 100 with a 100x/1.3 Fluar objective and a cooled MicroMax CCD camera (Princeton Instruments, Trenton, NJ) driven by a MetaView Imaging system (Universal Imaging Corporation), a Zeiss Axiovert Observer Z.1 (63x/1.43 objective, with Apoptome mode, camera Axiocam 503 mono) driven by ZEN software, or a Leica TCSSP confocal microscope. Time-lapse imaging was performed using either a monochromator (Polychrome II, TILL Photonics, Germany), a LED source (Colibri 7, Zeiss), or lasers (488 nm, 561 nm, 647 nm, Leica) for excitation.

In some experiments, cells were labelled with DChol or with the fatty acid probe Bodipy-C12-568. After a washing step, the fluorescent lipid droplets were counted. The images were thresholded, binarized and processed for particle counting using Image J software (NIH). In living cells, fluorescence imaging was performed with imaging buffer (10 mM Hepes, pH 7.4, 145 mM NaCl, 4.5 mM KCl, 1.2 mM MgCl_2_, 1.2 mM CaCl_2_) or in Hepes-buffered (10 mM, pH 7.4) culture medium (without phenol-red) with or without 10% FCS.

To study the effect of cycloheximide on the development of lipid droplets, HEK293 or HeLa cells cultivated in 3 cm glass-bottom dishes were incubated with cycloheximide (20 µM) for 30 min up to 12 h at 37°C. Thereafter, Bodipy 493/503 (4 µM) was added for 30 min. After a washing step, the cells were imaged and droplets were counted as described above.

To analyse the kinetic and reversibility of the translocation process of ACAT, HEK-ACAT cells were cultivated in a temperature controlled microchamber on the Leica setup. The translocation of ACAT was initiated by eflucimibe (0.5 µM). Images were captured each 20 sec. The vesiculation process was quantified by using the particle counting module of Image J (NIH). Images were background subtracted, stacked, and particles were counted. After washing-off eflucimibe the reversibility of the vesiculation was initiated by addition of BSA (10%) or BSA-bound oleic acid (400 µM OA in 10% BSA). Sigmaplot 8.0 was used to fit the data according to the appropriate functions and to draw the graphs. Colocalization of fluorescent structures was calculated using the Pearson correlation coefficient (Image J, NIH).

### Gel electrophoresis, Western blot and immunocytochemistry

In most cases, proteins of cellular membranes were separated by SDS-PAGE. Prior to gel electrophoresis the samples were heated for 10 min either at 40°C, 70°C or 95°C in SDS loading buffer. In some experiments, the non-denaturing Blue native gel electrophoresis was performed according to Schägger [54]. The proteins that have been separated by gel electrophoresis, were transferred to nitrocellulose and were subjected to immunoblot analysis. After a blocking step with i-Block for 1 h, the immunodetection of ACAT was performed using anti-ACAT/SOAT antibody (1:200 in i-Block) and anti-mouse IgGκ-BP (1:1000 in i-Block, HRP conjugated) as the secondary antibody. Alternatively, anti-mouse-IgG binding protein (HRP conjugated) was used as the antibody detecting reagent. Anti-VDAC (1:100) was used as a loading control. The immunoblot was developed using Western Blotting Luminol Reagent (sc-2048).

The cells that were processed for immunocytochemistry were grown on an 18 mm coverslip placed either in a 6 well plate or in a microchamber. The cells were fixed with 3.7% paraformaldehyde in PBS for 15 min at 25°C. After washing-off the fixation reagent, the cells were incubated with the appropriate antibody, i.e., anti-calnexin (mouse) (1:100), anti-Golgin (1:50), or anti-Sec31 (1:50). As the secondary antibody anti-mouse-Cy3 (1:1000 in PBS / 5% FCS) was applied. All antibodies were diluted in PBS / 5% FCS. The cells were either subsequently imaged in PBS or were mounted in Mowiol 4-88 before imaging.

### Crosslinking studies

AC29-ACAT cells were used to analyse the electrophoretic mobility of ACAT in dependence of ACAT inhibition and sucrose. The cells grown in 6-well plates were treated or not treated with sucrose or eflucimibe for 1 h. Then, the membrane-permeable aminoreactive crosslinker disuccinimidyl suberate (DSS) (dissolved in DMSO) was added to the cells at increasing concentrations (0.01 mM, 0.1 mM, 1 mM) for 30 min at 37°C. After washing-off the crosslinker with PBS, the cells were harvested by scraping and were immediately homogenized in PBS by sonication (7 sec, 40%, using HTU soni 130, Heinemann, Germany). After centrifugation (for 1 h at 32,000g) the homogenate pellet was resuspended in SDS loading buffer. Proteins were heated at 70°C for 10 min and separated by SDS-PAGE, followed by immunoblot analysis with anti-ACAT/SOAT antibody as described above.

### ACAT activity assays

In cells, the cholesterol esterification was measured using DChol as substrate. Cells grown on Petri dishes were labelled with DChol-MβCD (7 µM) or (as control) with Chol-MβCD (7 µM) in culture medium/10% FCS at 37°C in the absence or presence of an ACAT inhibitor (iACAT) (e.g., eflucimibe at 0.4 µg/ml). The medium was removed and the cells were washed twice with PBS. Then, the cells were harvested by scraping and their lipids were extracted as described above.

In microsomes, the assay was performed as follows. AC29-ACAT cells grown on 10 cm Petri dishes were washed with PBS, were then harvested by a cell scraper and centrifuged for 5 min at 500g. The cell pellet (∼1 x 10^7^ cells) was resuspended in 500 µl PBS, sonicated (for 4 s at 40% intensity using a Branson sonifier), and was then centrifuged for 1 h at 32000g. The pellets were frozen in aliquots at - 70°C. To measure the ACAT activity, the membrane pellet (2 x 10^6^ cells) was resuspended in 300 µl assay buffer (20 mM Hepes, pH 6.8, 2 M KCl). After addition of 1 µl DChol-MβCD (3 mM stock), the assay was mixed and incubated for 5 min at 25°C. The esterification reaction was initiated by addition of 1 µl oleoyl-CoA (final 0.8 – 64 µM from increasing stock solutions). After incubation for 2 h at 37°C the lipids were extracted, and were separated by TLC on silica gel plates using petrolether:diethylether:acetic acid (8:2:1, vol%) as the solvent system. DChol and DChol ester bands were quantified by fluorimetry (see above).

## Abbreviations

ACAT: acyl-CoA:cholesterol acyltransferase
BSA: bovine serum albumin
CHX: cycloheximide
DSS: disuccinimidyl suberate
CE: cholesterol ester
DChol: Dansyl-cholestanol
FC: free cholesterol
iACAT: ACAT inhibitor
INSIG: insulin-induced gene protein
MAM: mitochondria associated membrane
MβCD: methyl-β-cyclodextrin
MLX: Max-like protein X
OA: oleic acid
SREBP: sterol response element binding protein
SOAT: sterol O-acyltransferase
SCAP: SREBP cleavage-activating protein
TAG: triacylglycerol.

## Acknowledgements

We thank Larisa Cristea and Conny Trossen for their help in this research project. AC29 cells were kindly provided by T.Y. Chang (Dartmouth, U.S.A.).

## Author contributions

GG: design and planning of the study, acquirement and analysis of data, interpretation of the results, drafted the manuscript. LA and AS performed experiments, and analyzed the data. All authors critically reviewed iterations of the manuscript and approved the final draft for submission.

