## Supplemental Figures for "Reversible translocation of acyl-CoA:cholesterol acyltransferase (ACAT) between the endoplasmic reticulum and vesicular structures"

### SUPPLEMENTARY FIGURES

**Fig. S1**

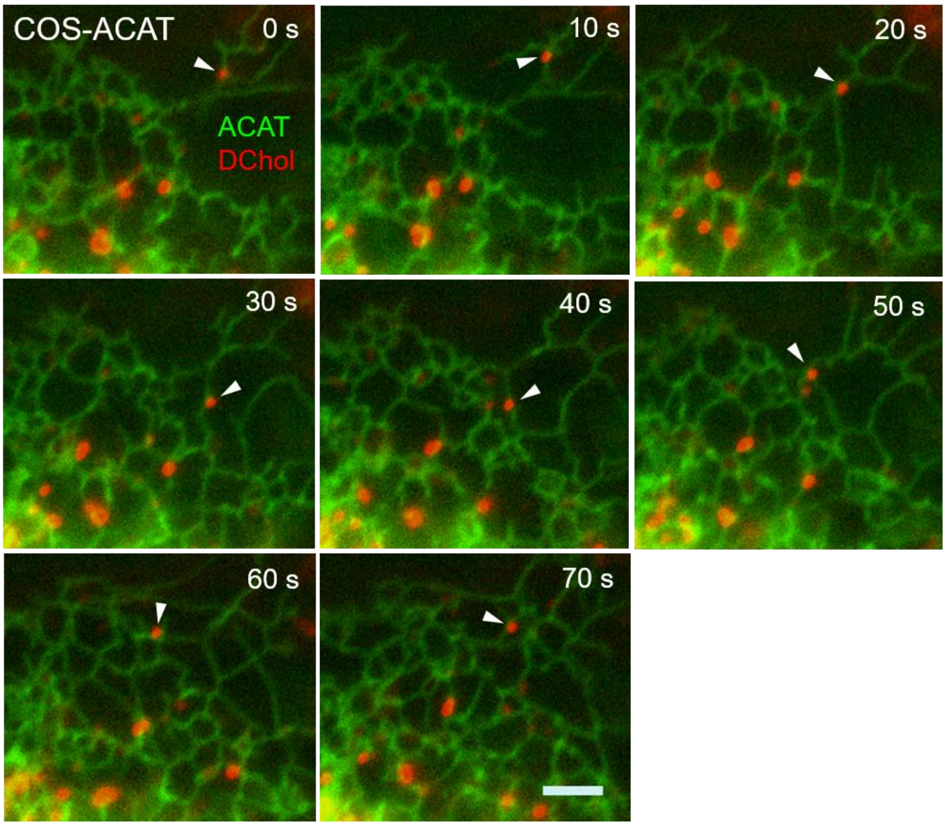


**Fig. S1: Most DChol containing droplets are associated with the endoplasmic reticulum.** COS-ACAT cells labeled with DChol-MβCD (10 µM) for 30 min were washed and further incubated for 7 h. The distribution of ACAT-GFP (green) and DChol/ester (red) was imaged. Most DChol containing droplets are closely associated with the ER as shown for the droplet marked with arrowhead (bar, 3 µm) (Image J processing, using flattened background).

**Fig. S2**

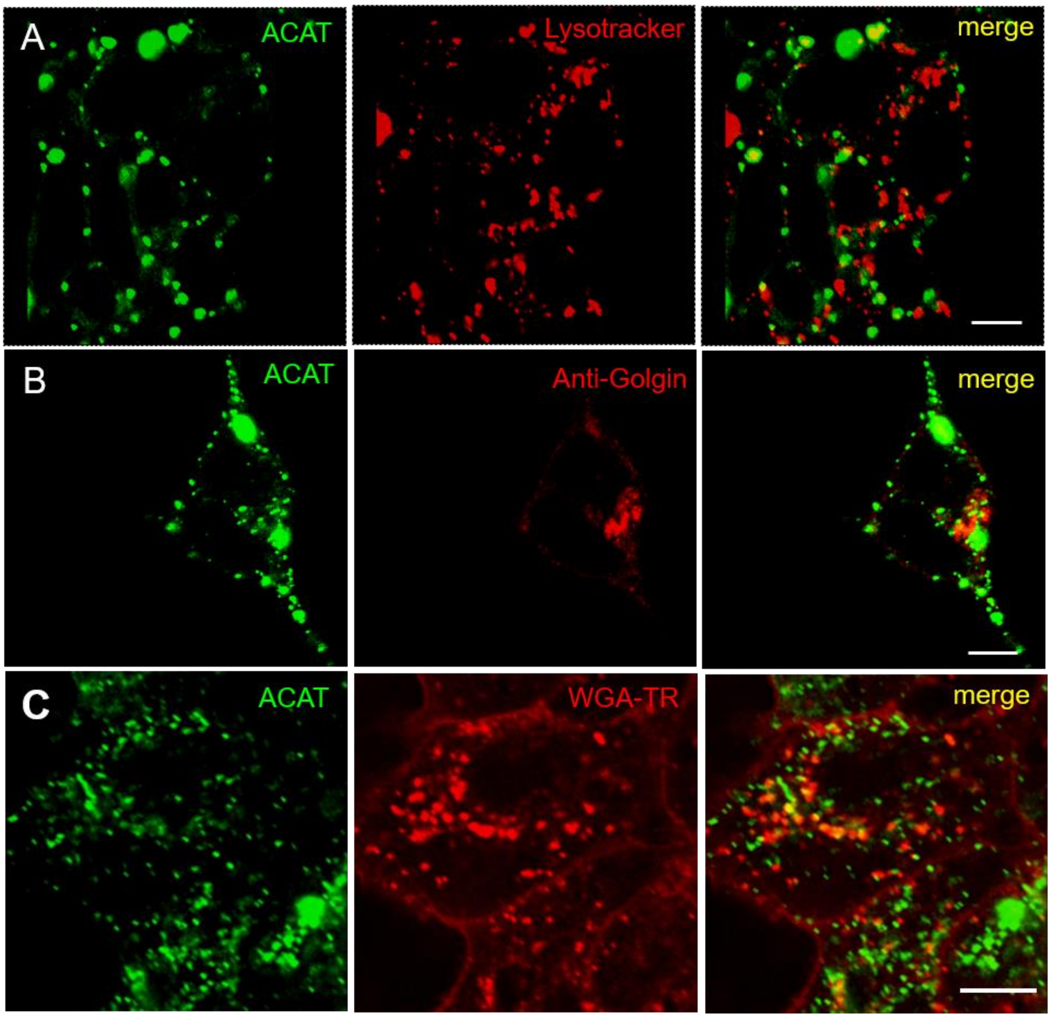


**Fig. S2: Colocalization of ACAT vesicles with organelle-specific markers.** HEK-ACAT cells were treated with eflucimibe for 1 h. Then, living cells were incubated with lysotracker Red (50 nM, 10 min) (A), with wheat germ agglutinin, Texas Red-X Conjugate (WGA-TR) (10 µg/ml, 10 min) (C), or left untreated (B). The free dye was washed off and images were captured. Formaldehyde-fixed and permeabilized cells were stained with anti-Golgin-97 (1 µg/ml) / anti-mouse-IgG-Cy3 (B) to stain the Golgi compartment (B). ACAT vesicles are shown by their green GFP fluorescence (A-C) (Bars, 10 µm).

**Fig. S3**

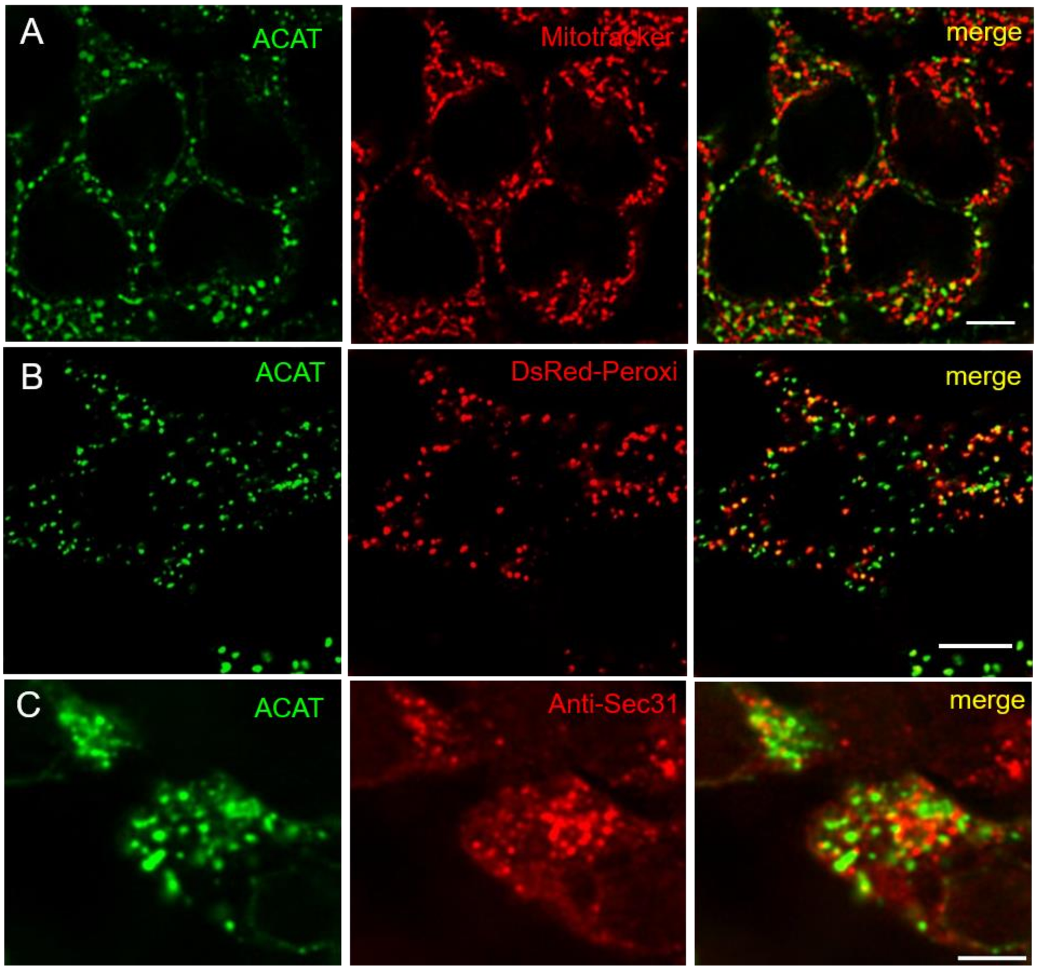


**Fig. S3: Colocalization of ACAT vesicles with organelle-specific markers.** HEK-ACAT cells (A, C) or HEK cells stably co-expressing encoding ACAT-GFP and DsRed-Peroxi (with ´SKL` targeting signal) (B) were treated with eflucimibe for 1 h. To visualize mitochondria, living cells were incubated with Mitotracker CMTMRos (200 nM, 10 min, 30°C) (A). The free dye was washed off and images were captured after fixation with formaldehyde. Peroxisomes were detected in formaldehyde-fixed HEK cells co-expressing ACAT-GFP and DsRed-Peroxi by capturing the fluorescence of DsRed (B). Formaldehyde-fixed and permeabilized HEK-ACAT cells were incubated with anti-sec31 / anti-mouse-IgG-Cy3 to stain ER exit sites (C). ACAT vesicles are shown by their green GFP fluorescence (A-C) (Bars: 10 µm in A and B, 5 µm in C).
